# Stoney vs. Histed: Quantifying the Spatial Effects of Intracortical Microstimulation

**DOI:** 10.1101/2021.08.12.456091

**Authors:** Karthik Kumaravelu, Joseph Sombeck, Lee E. Miller, Sliman J. Bensmaia, Warren M. Grill

## Abstract

**Background:** Intracortical microstimulation (ICMS) is used to map neural circuits and restore lost sensory modalities such as vision, hearing, and somatosensation. The spatial effects of ICMS remain controversial: Stoney and colleagues proposed that the volume of somatic activation increased with stimulation intensity, while Histed et al. suggested activation density, but not somatic activation volume, increases with stimulation intensity.

**Objective:** We used computational modeling to quantify the spatial effects of ICMS intensity and unify the apparently paradoxical findings of Histed and Stoney.

**Methods:** We implemented a biophysically-based computational model of a cortical column comprising neurons with realistic morphology and representative synapses. We quantified the spatial effects of single pulse ICMS, including the radial distance to activated neurons and the density of activated neurons as a function of stimulation intensity.

**Results:** At all amplitudes, the dominant mode of somatic activation was by antidromic propagation to the soma following axonal activation, rather than via trans-synaptic activation. There were no occurrences of direct activation of somata or dendrites. The volume over which antidromic action potentials were initiated grew with stimulation amplitude, while the volume of somatic activations did not. However, the density of somatic activation within the activated volume increased with stimulation amplitude.

**Conclusions:** The results resolve the apparent paradox between Stoney and Histed’s results by demonstrating that the volume over which action potentials are initiated grows with ICMS amplitude, consistent with Stoney. However, the volume occupied by the activated somata remains approximately constant, while the density of activated neurons within that volume increase, consistent with Histed.

**HIGHLIGHTS:**

- Implemented a biophysically-based computational model of cortical column comprising cortical neurons with realistic morphology and representative synapses.
- Quantified the spatial patterns of neural activation by intracortical microstimulation to resolve the paradoxical findings of Stoney et al., 1968 and Histed et al., 2009.
- The dominant mode of neural activation near the electrode was direct (i.e., via antidromic propagation from direct activation of the axon) and not trans-synaptic.
- The dominant effect of increased ICMS intensity was to increase the density of activated neurons but not the volume of activation.

## INTRODUCTION

Intracortical microstimulation (ICMS) provides a causal means to interrogate neural circuits and their role in behavior. In a classic study, ICMS applied to medial temporal cortex (MT) systematically biased visual motion perception in a manner consistent with visually induced firing rates, confirming the role of MT in visual motion processing (*1*). This strategy has been employed in a variety of contexts to implicate specific populations of neurons in specific behaviors, from perception to movement to cognition (*2–16*). The success of ICMS in eliciting sensory percepts led to its application as a means to restore lost sensation (*17, 18*). For example, ICMS delivered to somatosensory cortex elicits touch sensations experienced on the otherwise insensate hand (*17, 19*), and the restoration of touch sensations improves the dexterity of brain-controlled bionic hands (*20*).

Despite the widespread use of ICMS, the neuronal correlates thereof are poorly understood, in part because ICMS-induced electrical artifacts obscure ICMS-evoked neuronal activity (*21–25*). Even at a coarse level, the distribution of neuronal activity around the stimulating electrode is not well understood. According to one theory, the threshold to activate a neuron is proportional to the square of the distance to the stimulating electrode and increasing stimulation intensity activates a growing spherical volume of neurons (*23*). Consistent with this hypothesis, the sensory or cognitive consequences of ICMS through a given electrode can be at least roughly predicted from the properties of neurons recorded from that electrode, as in the experiment described above. However, axons are activated at lower currents than somata (*26–30*) so the spatial pattern of activation should be determined not by the neighboring cell bodies but rather by the distribution of axons passing through the spherical volume. Consistent with this hypothesis, optical imaging of ICMS-induced activity revealed a more broadly distributed and idiosyncratic spatial pattern of activation (*31*).

To resolve this paradox, we implemented a biophysically-based computational model of cortical neurons that incorporated realistic morphologies and membrane properties. First, we confirmed that axons were more sensitive to electrical stimulation than were cell bodies, and we observed that the volume over which axons were activated grew with stimulation amplitude, as expected. Next, we replicated the findings that the dominant mode of somatic activation was direct (i.e., antidromic propagation subsequent to axonal activation) rather than indirect (i.e., trans-synaptic activation) at all ICMS amplitudes, and that activation of axons around the stimulating electrode resulted in somatic activations distributed over a wide volume of tissue. Surprisingly, the volume over which cell bodies were activated was nearly independent of stimulation amplitude. However, the density of activated cell bodies was proportional to the ICMS amplitude and dropped rapidly as a function of distance from the stimulating electrode. In conclusion, ICMS activates cell bodies within a volume that is relatively insensitive to amplitude. However, the density of somatic activation within this volume is dependent on ICMS amplitude. These findings reconcile two seemingly incongruent hypotheses about the neuronal correlates of ICMS.

## METHODS

### Overview

The computational model comprised three parts: (1) single-cell cortical neurons with realistic axon morphology, (2) representative synapses, (3) single-cell cortical neurons with representative synapses arranged in a cortical laminar format with dimensions derived from histological sections from the rhesus macaque.

### Morphology of cortical neurons

We implemented a computational model of a population of biophysically-based Hodgkin-Huxley style multi-compartment cortical neurons, adapted from the Blue Brain cell library (*32, 33*). The model neurons included five different cortical cell types: Layer 1 neurogliaform with dense axonal arbor (L1 NGC-DA), Layer 2/3 pyramidal cell (L2/3 PC), Layer 4 large basket cell (L4 LBC), Layer 5 thick tufted pyramidal cell (L5 TTPC) and Layer 6 tufted pyramidal cell (L6 TPC) (Fig 1A). Further morphological diversity was achieved by introducing random variations to branch length and angle to create clones of each cell type (Fig S1) that had the same ion channel properties as the standard model but different morphologies (*32*). Specifically, each branch length in the standard model was scaled by a random number sampled from a Gaussian distribution of mean 0% and standard deviation 20%. Further, each of the two subtrees at a bifurcation point was rotated by an angle in degrees, sampled from a Gaussian distribution of mean 0° and standard deviation 10°. The rotation happened around the vector determined by the point at the base of the subtree and the first principal component of all arbor lengths in the subtree.

**Figure 1:**
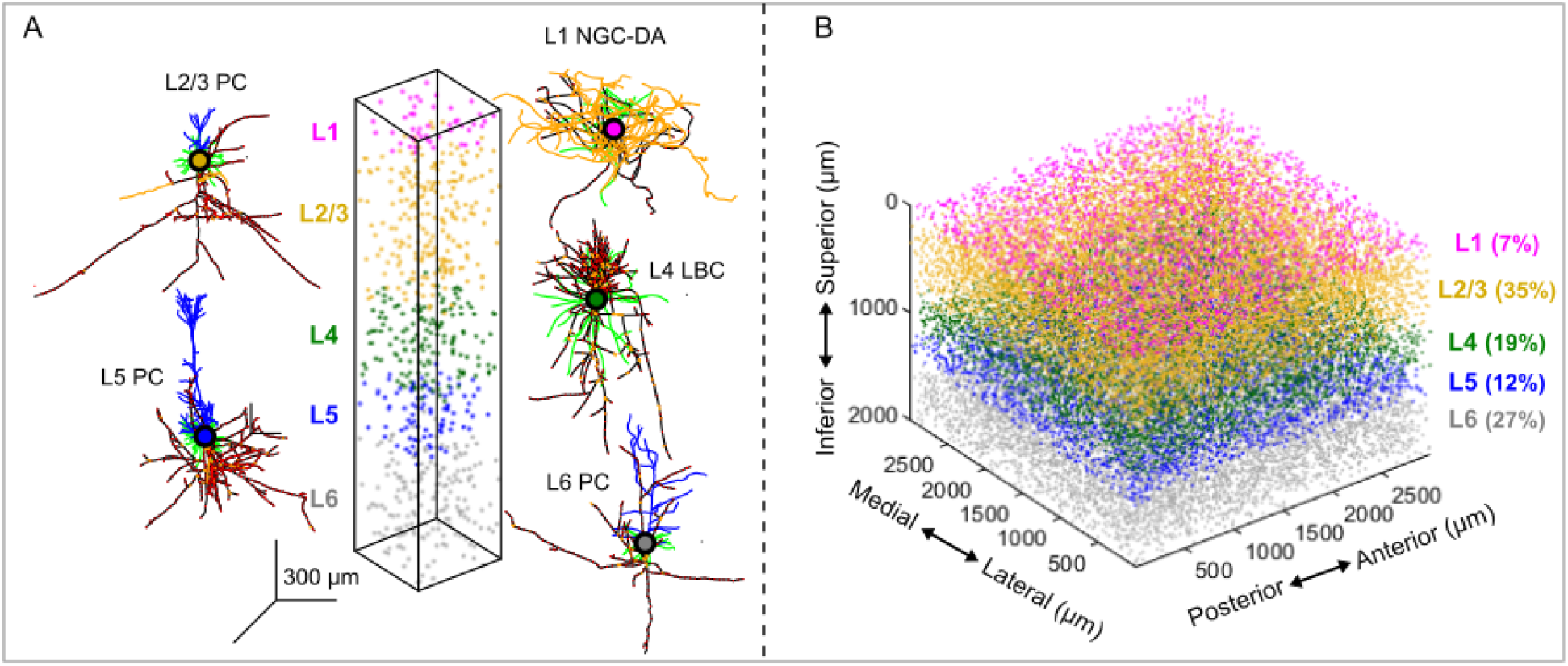
Biophysically-based computational model of somatosensory cortex. (A) 3D morphology of Layer (L)-1 neuroglial cell, L2/3 pyramidal cell (PC), L4 basket cell, L5 thick tufted (TT) PC, L6 thick (T) PC. Model neurons were adapted from the Blue Brain library and had realistic axon morphologies (*32, 33*). Blue – apical dendrites, green – basal dendrites, yellow – unmyelinated axon, black – myelin nodes, filled circle – soma. Model axons were myelinated, and the diameter range was consistent with those found in rhesus macaques (*33, 34*). Cortical neurons were arranged in a column comprising five layers with a dimension of 400 μm x 400 μm x 2000 μm. Cortical thickness was determined from histological sections of somatosensory cortex in rhesus macaque (*37*). (B) The cortical column shown in (A) was arranged in an 8*8 grid with a dimension of 3000 μm x 3000 μm x 2000 μm. The relative thickness of each layer was based on estimates from the visual cortex of rhesus macaque (*38*) and is indicated in parenthesis. A uniform density of 2000 neurons/ was used across all layers, resulting in a total cell count of 36,000 neurons for the slab model with volume 18.

The cortical neurons’ axons were inappropriately unmyelinated in the original Blue Brain publication and were unexcitable at low ICMS intensities (Fig S2A). Aberra et al., fixed the excitability issue by myelinating the axons (*33*), making them excitable with low ICMS intensities (Fig S2B). We adapted the myelinated version of model neurons for our study (ModelDB accession number: **241165**). The distributions of diameters across all axonal compartments (node of Ranvier, myelin, and unmyelinated segments) are shown in the original publication and matched those in rhesus macaques (*33, 34*). Further details on the parameters for myelination (such as *g* ratio, 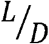 ratio) can be found in the original publication (*33*). The model neurons included 13 known classes of Hodgkin-Huxley-type ion channels across soma, dendrite and axon: D-type potassium, A-type potassium (Kv3.1), stochastic potassium, transient potassium, persistent potassium, transient sodium (soma), transient sodium (axon), persistent sodium, low-voltage-activated calcium, high-voltage-activated calcium, M-current, H-current, and calcium-activated potassium. Parameters such as conductance density, reversal potential, and time constant for the different ion channel models can be found in the original publication (*32*).

### Representative synaptic connections

Although it would be desirable to simulate the complete cortical microcircuitry, this is computationally prohibitive, so to study synaptic effects of ICMS, we incorporated only representative synapses. For each cell type, the location of postsynaptic compartments (across the dendritic arbor) where an incoming synaptic connection is received were mapped in the Blue Brain project (*32*). Fig 2A shows the locations of excitatory and inhibitory postsynaptic connections received by L5 PC cells from five different cell types (L1 NGC-DA, L2/3 PC, L4 LBC, L5 PC and L6 PC). We show synaptic inputs from only five cell types in Fig 2, whereas in the model each cell type received synaptic inputs from 55 different cell types. All input types were cortical in origin and did not include connections from thalamus, subcortex, etc. Fig 2B shows the Euclidean distance of each postsynaptic compartment from the soma for an L5 PC.

**Figure 2:**
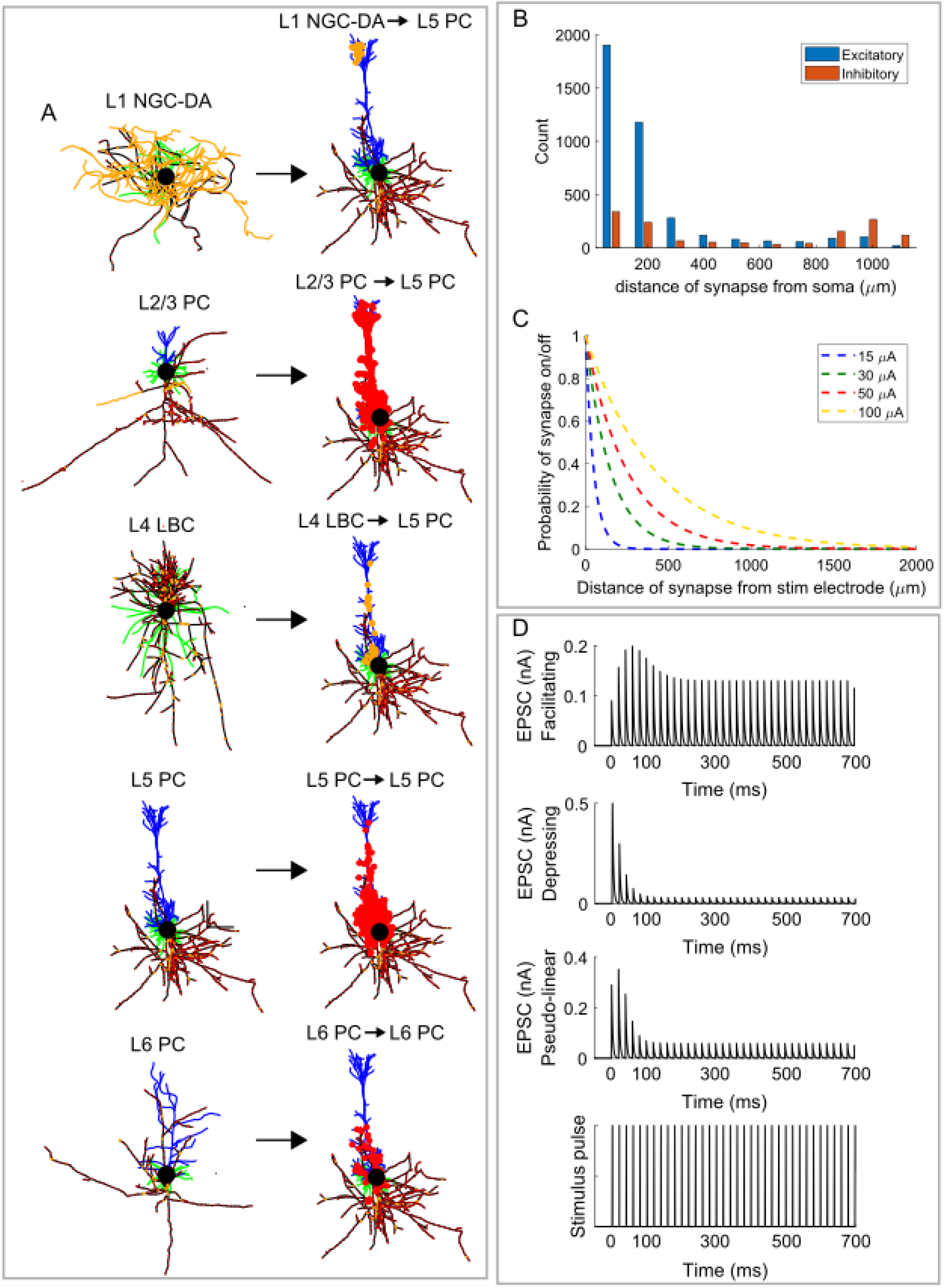
Properties of the representative synapses. (A) Synaptic inputs to L5 PC from L1 NGC-DA, L2/3 PC, L4 LBC, L5 PC and L6 PC. Each cortical neuron (L1 NGC-DA, L2/3 PC, L4 LBC, L5 PC and L6 PC) received synaptic inputs from 55 different presynaptic sources. All sources were cortical and did not include inputs from other structures such as the thalamus, subcortex, etc. Excitatory and inhibitory postsynaptic compartments are shown in red- and yellow-filled circles, respectively. Inhibitory synapse comprised GABA-A+GABA-B receptors and excitatory synapse were composed of AMPA+NMDA receptors. (B) The distribution of Euclidean distance of synapses from the soma of an L5 TTPC. (C) The probability of synaptic activation by extracellular stimulation was an exponential function of its distance from the electrode tip. Space constant for different stimulation amplitudes: 15 μA – 50 μm, 30 μA – 150 μm, 50 μA – 250 μm, 100 μA – 420 μm. (D) EPSC of AMPA synapse showing dynamics of facilitating, depressing and pseudo-linear synapse in response to 50 Hz stimulation.

For excitatory connections, each postsynaptic compartment included AMPA and NMDA receptor kinetics. Similarly, inhibitory connections included both GABA-A and GABA-B receptor kinetics. The parameterization of unitary synaptic connections was explained in detail in the original publication (*32*). Experimental postsynaptic responses were fit using a stochastic version of the Tsodyks-Markram model for dynamic synaptic transmission including short-term facilitation (STF) and depression (STD) effects described by the following differential equations:

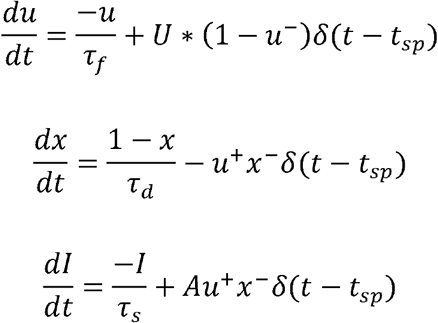

STD is simulated through variable *x*, representing the fraction of resources available after neurotransmitter depletion. Variable *u* denotes the fraction of resource ready to use for an upcoming spike and is used to simulate STF. *τ_f_* is the decay time constant of the variable *u* and *τ_d_* is the recovery time constant of variable *x. u*, representing synaptic efficacy, is the fraction of recovered neurotransmitter that is instantaneously released at each incoming spike. *I* represents the postsynaptic current generated by a spike arriving at the time *t_sp_. τ_s_* is the decay time constant of the variable *I* and *A* is the maximum synaptic response possible following release of all neurotransmitters. The interplay between parameters *τ_f_, τ_d_* and *U* is used to generate the effects of depression and facilitation. These parameters can be obtained from the original publication (*32*). Fig-2D shows example synaptic dynamics (EPSC for AMPA synapse) to a 50 Hz stimulating train. Finally, values of synaptic conductance across all connections received by a given cell type can be found in the original publication, obtained by fitting the amplitude of *in silico* somatic postsynaptic potential responses to the *in vitro* responses (*32*).

The probability of activation of a synapse by extracellular stimulation was an exponential function of its distance from the electrode tip. The space constants of the exponential curves were derived as follows. First, model neurons for each cell type were placed in a spherical volume of radius 2000 μm. Second, stimulation at different intensities was applied at the center of the volume, and the location of axons where action potentials were initiated was determined. Third, the number of activated axons as a function of the distance from the electrode tip was fit by an exponential curve, and the space constants were calculated. Finally, the median space constant across the five cell types was used to compute the probability of synaptic activation. Fig 2C shows the probability density function of the exponential distribution used to determine synaptic activation by stimulation. Space constants were larger for higher stimulation amplitudes indicating that larger amplitudes can activate pre-synaptic terminals farther from the electrode tip. This method of estimating space constants for synaptic activation is justifiable because axon terminals (presynaptic terminals) are the most sensitive axonal element activated by extracellular stimulation (*29, 33, 35, 36*). Although other geometric discontinuities such as axonal branches can be activated by stimulation, their proportion (20%) is low compared to axon terminals (80%) (*33*). For each synapse, the delay between stimulation pulse and synapse being turned on was chosen from a uniform distribution between 0 to 2ms. This delay was randomized for each synapse to account for axonal propagation time and synaptic delay.

### Cortical column model

The cortical neurons with representative synapses were arranged in a column with five layers (Fig 1A). The cortical thickness was derived from histological slices from the postcentral gyrus of rhesus macaques (*37*). We used a cortical thickness of 2000 μm for our cortical column like those seen in areas 1 and 2, with overall dimensions of 400 μm x 400 μm x 2000 μm (Fig 1A). The single cortical column model was extended to 64 columns (slab model) placed in an 8*8 grid with dimensions 3000 μm x 3000 μm x 2000 μm (Fig. 1B). Each layer’s relative thickness in the column was based on estimates from the primary visual cortex of rhesus macaque (*38*): L1-7%, L2/3-35%, L4-19%, L5-12% and L6-27%. The actual cell density in somatosensory cortex of macaque is in the order of ~100,000 neurons/*mm*^3^ (*39*). However, we used a density of 2000 neurons/*mm*^3^ to save computational time. This resulted in 36,000 neurons for the slab model in a volume of 18 *mm*^3^. Further, the density of 2000 neurons/*mm*^3^ was uniform across the five cortical layers. The cell count per layer was: L1-2520, L2/3-12600, L4-6840, L5-4320 and L6-9720. For each cell type, an equal proportion (20%) of cells from each of the five clones was used. For example, in L1, 20% of 2520 (504) cells were L1 NGC-DA clone-1, 20% of 2520 (504) cells were L1 NGC-DA clone-2, and so on. Cells were positioned randomly within a layer based on a uniform distribution with upper and lower bounds derived using the lamina dimensions mentioned above. Further, each neuron was randomly rotated around its somatodendritic axis at an angle obtained from a uniform distribution between 0° to 360°.

### Parameters for ICMS simulation

The stimulating tip of the electrode was approximated as a point current source (*40, 41*). Extracellular potential *V_e_*(*i,j*) due to the point current source in each compartment (*j*) of each model cortical neuron (*i*) was computed using the equation,

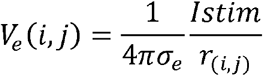

where *Istim* is the stimulating current, *r_i,j_* is the distance from the stimulating electrode tip to each compartment of the model cortical neuron, and *σ_e_* is the isotropic and homogeneous conductivity of the extracellular medium (0.3 S/m) (*42*). All simulations involved stimulation by a single biphasic pulse (cathodic first) with a fixed width of 200 μs/phase and an interphase interval of 53 μs. We tested clinically relevant stimulus intensities ranging from 5 to 50 μA in steps of 5 μA (*14, 17*). All neurons that had a neural element (dendritic, somatic or axonal compartments) within 15 μm from the electrode tip were excluded from the analyses since *V_e_* estimation using a point source would be inaccurate for such close distances without considering the 3D geometry of the microelectrode (*40*). Further, to quantify the proportion of direct versus indirect somatic activation by ICMS, we ran simulations in the model with and without synapses. Synapses were turned off by setting the synaptic conductance to zero (i.e., *g_syn_* = 0 *nS*). We consider the number of directly activated neurons to be the number of somas activated in the model without synapses. For quantifying indirect activation, we calculated the difference in total somatic activation between the models with and without synapses. The proportion of directly vs. indirectly activated neurons was quantified using a 13 pulse ICMS train at 2 Hz and 300 Hz, as higher frequencies are expected to activate more neurons indirectly by postsynaptic temporal summation (*43*).

Simulations were implemented in NEURON 7.7 with equations solved using the backward Euler method with a time step of 0.025 ms (*44*). The values of *V_e_* were coupled to each neuronal compartment using the *e_extracellular* mechanism (*44*). The simulation was parallelized by distributing the total number of neurons (36,000) across 400 processors in a round-robin fashion (90 neurons/processor) (*45*).

### Statistical Analysis

The following statistical tests were undertaken to infer effects of ICMS intensity on axonal and somatic activation distances. First, we tested the assumption of normality of activation distance at each ICMS intensity using the one-sample Kolmogorov-Smirnov test. Activation distance data were not normally distributed. Therefore, statistical inferences were made on the effect of ICMS intensity on activation distance using the Kruskal-Wallis one-way analysis of variance (ANOVA). When the omnibus test statistic revealed significance at *p<0.05*, we performed Dunn-Sidak’s test for *post hoc* paired comparisons between individual ICMS intensities (*46*). All statistical analyses were performed using MATLAB (Mathworks, Natick, MA).

## RESULTS

### Effect of ICMS intensity on axonal vs. somatic activation distance and density

ICMS at different intensities was delivered through a stimulating electrode was positioned in L4 of the cortex. For each soma activated by ICMS, we tracked the location on the axon where the action potential (AP) was initiated by locating the compartment with the earliest latency. Fig 3A shows the location of AP initiation in the axon for each of the activated somas shown in Fig 3B at intensities 5, 15, 30 and 50 μA. The AP was always initiated in the axon, predominantly at a geometric discontinuity such as a branch or termination, and there were no instances where an AP was initiated in the soma or dendrite. The somatic activation was due to antidromic propagation of the AP from an axon activated close to the stimulation electrode. We collapsed the 3D column data shown in Fig 3A, B to the (2D) sagittal plane, where the radial coordinate denotes the distance of the activated soma or axon from the electrode, and the angular coordinate of each neuron was randomly drawn from a uniform distribution between 0° to 360° to spread out the data (Fig 3C, D).

**Figure 3:**
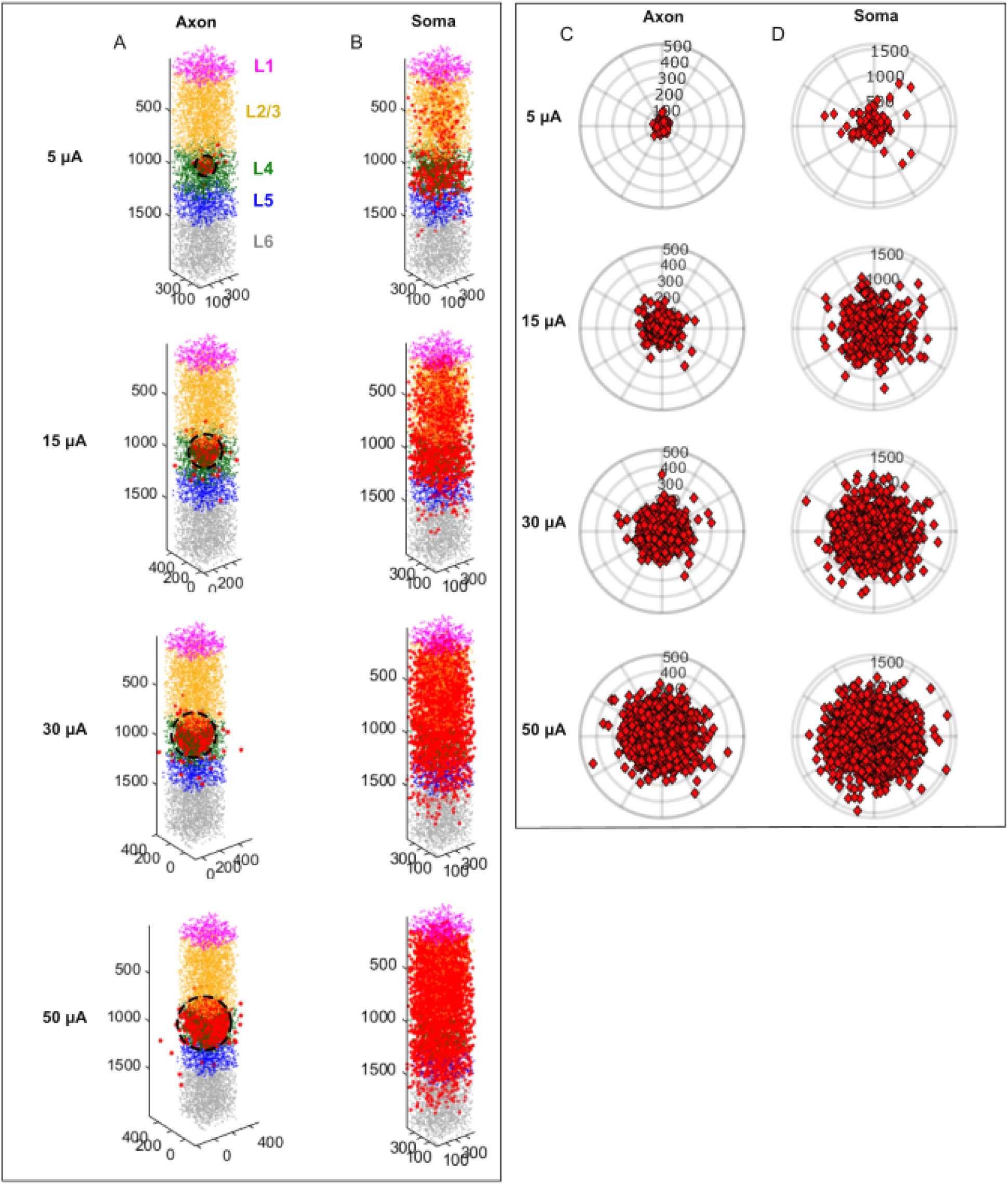
Response to intracortical microstimulation (ICMS) at different intensities. (A) Axonal locations where ICMS initiated the action potential (red dots). Following initiation in the axon, action potential propagated antidromically back to soma. (B) Location of activated soma (red dots) to ICMS. Activation of axons close to the electrode tip (shown in A) activated somas both local and distant from the stimulation electrode. 2D polar plane (sagittal) was used to visualize the location of activated somas and axons shown in (A) and (B). Polar plots (in sagittal plane) showing the location (radial distance) of activated axon (C) and soma (D). Theta for each neuron was randomly assigned from a uniform distribution between 0-360 degrees.

The spread of activated axons increased as the stimulation intensity increased, while the spread of activated somata did not. Indeed, the radial distance of axonal activation increased with stimulation intensity (Fig 4A) and stimulus intensity exerted a significant effect on activation distance (*p* ~ 0, *Kruskal – Wallis ANOVA, χ*^2^ (9) = 1530). Post-hoc analysis revealed a significant difference in activation distance across stimulation intensities (Fig 4A, *p* < 0.05, *Dunn – Sidak method*). In contrast, somatic activation distance increased only marginally with stimulation intensity (Fig 4A), though this effect was significant (*p* ~ 0, *Kruskal-Wallis ANOVA, χ*^2^ (9) = 205). Post-hoc analysis revealed a difference in activation distance between low (< 20 *μA*) and high (> 20 *μA*) stimulation intensities (Fig 4A, *p* < 0.05, *Dunn – Sidak method*). While the median radial distance for axonal activation increased fourfold from ~26 μm at 5 μA to ~105 μm at 50 μA, that for somatic activation increased less than two-fold from ~294 μm to ~455 μm over the same range of intensities (Fig-4A). Further, we compared the radial distance from the model with the radial distance predicted by Stoney’s 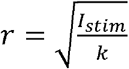 relationship for three different k (current-distance constants) from the literature (*47*): 300 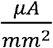, 1292 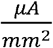 and 27000 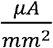 (Fig 4A). Stoney’s relationship was predictive of the increase in radial distance with stimulation intensity for axonal activation but not for somatic activation (Fig 4A).

**Figure 4:**
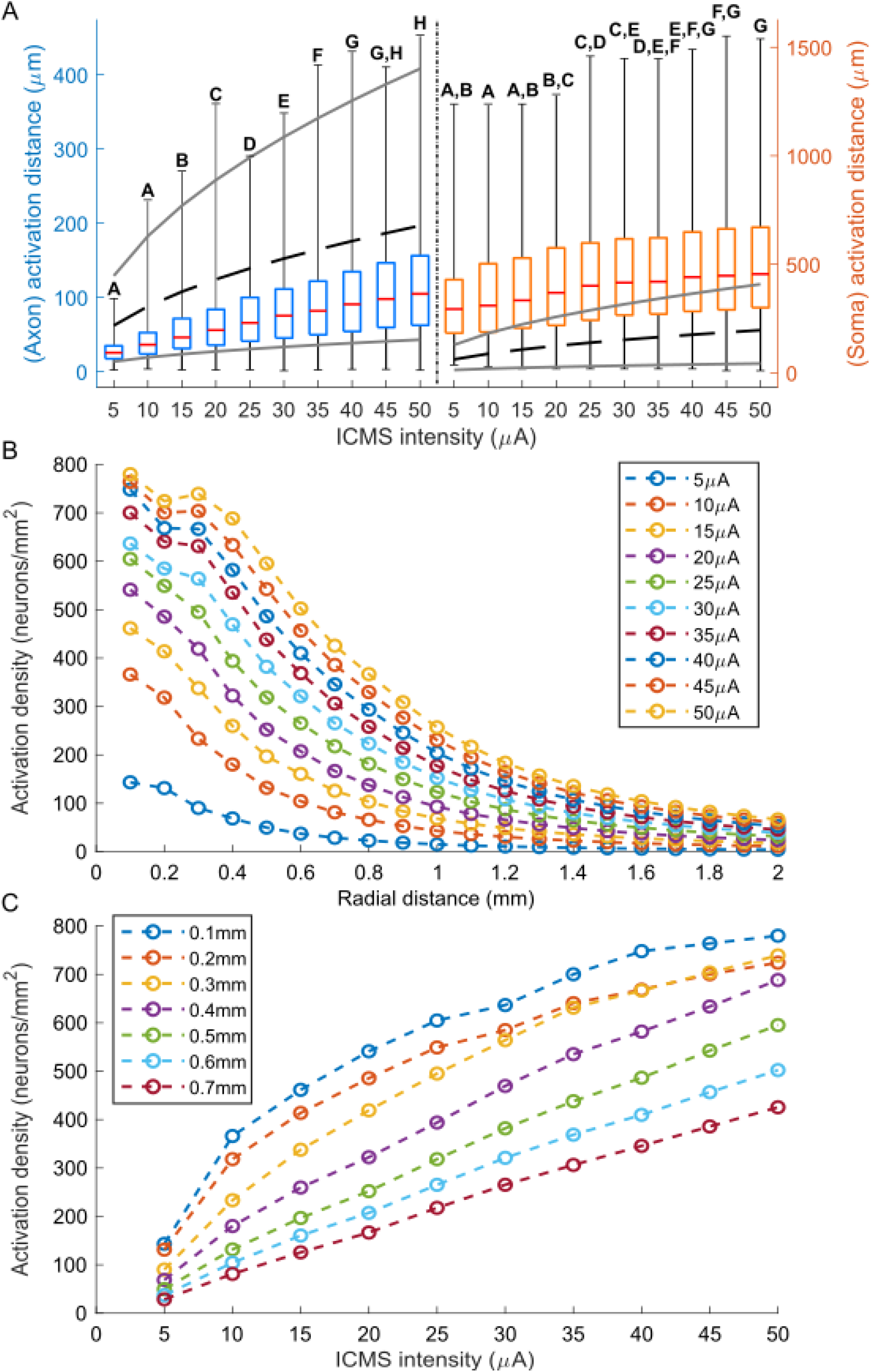
Quantification of response to intracortical microstimulation (ICMS) at different intensities. (A) Distribution of activation distance across somas/axons as a function of stimulation intensity. Activation (radial) distance was derived from the polar plots shown in Fig 3C, D. On each box, the central mark indicates the median, and the bottom and top edges of the box indicate the 25th and 75th percentiles, respectively. Whiskers extend to the absolute minimum and maximum. Kruskal-Wallis ANOVA identified effects of stimulation intensity on activation distance (p ~ 0). Stimulation intensities that do not share the same letter are significantly different (p<0.05, Dunn-Sidak method). The grey and black lines are the predicted activation distances based on Stoney’s 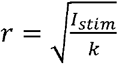 relationship for three different ‘k’ values: (1) 300 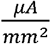 - upper gray trace, (2) 1292 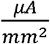 – middle black trace, (3) 27000 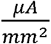 – bottom gray trace (*47*). (B) Somatic activation density as a function of radial distance for various stimulation intensities. (C) Activation density as a function of ICMS intensity for an area with different radii. Somatic activation density increased with stimulation intensity.

Since the somatic activation volume did not increase substantially with stimulation intensity, we tested whether the density of activated somata increased with intensity. The activation density decreased with distance from the electrode tip (Fig 4B), implying that the bulk of the somata activated by stimulation were near (< 400 μm) the stimulation electrode. The somatic activation density increased with stimulation intensity, particularly within 400 μm of the stimulus electrode tip (Fig 4C). The relationships between stimulation intensity and activation distance and activation density also held when expressed over the 2D axial plane (Fig S3).

### Effect of ICMS intensity on the proportion of direct vs. indirectly activated neurons

We determined whether somata activated by stimulation were directly activated (*via* antidromic propagation from directly activated axon) or activated indirectly (*via* activation of synapses) by running simulations with and without synapses (Fig 5 A, B). We quantified the number of neurons activated by ICMS at a low (2 Hz) and a high (300 Hz) frequency (Fig 5C2). We found that most neurons were activated directly rather than trans-synaptically and, although stimulation at a higher frequency led to a higher proportion of trans-synaptically activated neurons, indirect activation still accounted only for ~20% of the overall activation in this condition (Fig 5C1).

**Figure 5:**
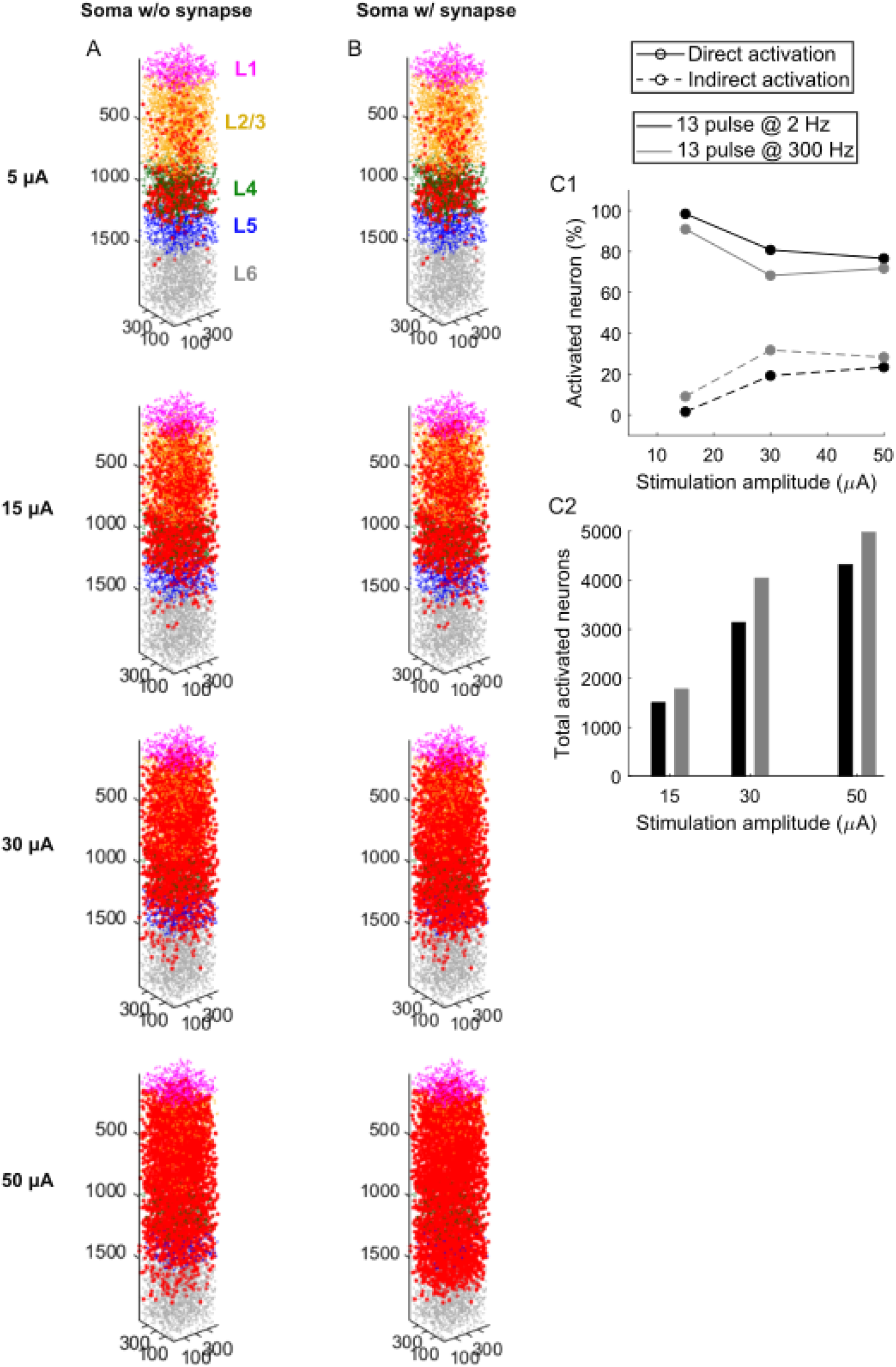
Proportion of spikes evoked by direct vs. indirect (synaptic) activation by ICMS at different intensities. A direct spike in a soma was generated antidromically due to direct activation of its axon. An indirect spike was trans-synaptically activated following ICMS mediated activation of synaptic inputs. Comparison of somatic activation between models (A) without and (B) with synapses for ICMS. (C) The proportion of direct vs. indirect spikes evoked by ICMS at different intensities. Most spikes were directly evoked across all stimulation intensities.

## DISCUSSION

We implemented a computational model of a cortical column comprising neurons with realistic morphology and representative synapses to study the spatial effects of ICMS and to resolve the seemingly paradoxical findings of Stoney et al. and Histed et al. (Fig. S4). First, the dominant mode of somatic activation was direct (i.e., antidromic propagation following direct axonal activation) rather than indirect (i.e., trans-synaptic activation) at all ICMS amplitudes. Second, the volume of axonal activation (i.e., location where action potential was initiated in the axon) grew with stimulation amplitude around the electrode tip, consistent with Stoney’s findings. Third, the volume of somatic activation did not increase with stimulation amplitude, consistent with the findings of Histed. Finally, somatic activation density within the activated volume increased with stimulation intensity.

### Neural (spatial) effects of ICMS intensity

Stoney and colleagues (*23*) used electrical stimulation and recording to measure the spread of ICMS induced activity, and they fit their strength-distance data using the following equation,

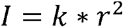

Where *I* is the stimulating current (in μA), *r* is the distance between the stimulating electrode and neuron (i.e., recording electrode) (in μm) and *k* is the current-distance constant (in *μA*/*μm*^2^). The value of *k* was estimated to be 1,292 *μA/mm*^2^. Accordingly, a stimulating current of 5 μA was predicted to activate somas within a radius of 63 μm, 10 μA within 88 μm, 20 μA within 124 μm, and so on. Our simulation results paint a more nuanced picture. First, very low stimulation currents (5 μA) resulted in a larger, nearly 300 μm radius of activation. Second, a transition from low (5 μA) to high (20 μA) currents only modestly increased the radius, from 294 μm to 369 μm. This suggests that Stoney and colleagues overestimated the increase in the somatic activation radius with ICMS intensity. However, the radial distance of axonal activation in our model was 26 μm at 5 μA and increased (doubled) to 56 μm at 20 μA. This increase in radial distance of axonal activation with stimulation intensity is consistent with Stoney’s 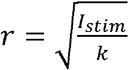 relationship.

Histed and colleagues (*31*) used two-photon imaging of calcium concentration to quantify the effect of ICMS intensity near the electrode tip in the superficial layer (L2/3) of both the rodent and cat visual cortex. They found somatic activation as far as 250 μm at currents as low as 9 μA. Next, displacing the stimulating electrode tip by 15 μm substantially modified the pattern of activated neurons. Further, they found no difference in activation pattern after blocking excitatory synaptic transmission and concluded that activation of axonal processes close to the electrode tip resulted in somatic activation (*via* antidromic propagation) both local and distant to the stimulation electrode even at low ICMS intensities. Stimulating at 5 μA resulted in a somatic activation distance of 294 μm in our model due to the activation of axons very close to the electrode tip and consistent with Histed’s findings.

Although Histed and colleagues did not systematically evaluate the effect of ICMS intensity on somatic activation distance, they concluded that the radial distance of somatic activation did not change substantially with increasing intensity. We conducted a systematic analysis of the effect of ICMS intensity on radial distance of somatic activation and activation density. The radial distance to the farthest activated soma did not increase substantially with ICMS intensity, whereas the activation density (i.e., the number of activated somas within the activated volume) increased with intensity.

Histed and colleagues observed a similar number and spatial distribution of activated somas before and after pharmacological blockade of excitatory synaptic transmission and concluded that activation was direct rather than trans-synaptic. Our finding that the proportion of neurons activated directly was much higher than indirectly activated neurons across all stimulation intensities (5-50 μA) was entirely consistent with their finding.

### Behavioral effects of ICMS intensity

ICMS of the somatosensory cortex is increasingly used to restore lost sensations such as touch and pressure (*11, 17, 19, 48, 49*). Two observations of the perceptual effects of ICMS are consistent with our modeling results. First, ICMS-induced cutaneous percepts are focal (*15, 17, 50*), often spanning a single finger pad or palmar whorl. These focal percepts suggest that ICMS induced neural activation is focal and close to the electrode, enabled by the underlying somatotopic organization. The observation that ICMS produces effects that can be predicted from the properties of local neurons has been observed in a variety of other contexts (*51*). In the model, somatic activation was focal to the extent that the distance to the farthest activated neuron did not change substantially with stimulation intensity. Second, the perceived magnitude of ICMS-evoked percepts increases linearly with stimulation intensity (*15, 17*), consistent with our observation of increased somatic activation density around the electrode tip with increasing stimulation intensity.

### Model limitations

Our cortical column model comprising neurons with realistic morphology and representative synapses is the most advanced modeling work to date to study the spatial effects of ICMS intensity, but it has several important limitations. First, the properties of ion channels used at the nodes of Ranvier and axon terminations are the same as those used at the initial segment. The ion channel properties at the initial segment were fit to experimental data in the Blue Brain study (*32*). However, fitting to electrophysiological data was not done for the other axonal compartments since they were not part of the final cortical column simulations in the Blue Brain project. Axonal ion channel properties become important during extracellular stimulation since, within axons, geometric discontinuities such as terminations and branches typically have the lowest threshold for extracellular stimulation (*30, 33, 52*).

Our model included only representative synapses and not full cortical connectivity between neurons. The model likely underestimates the proportion of indirectly activated neurons and thereby the spatial spread to ICMS because of the lack of full connectivity. Further, although we know that trans-synaptic activation can occur over widespread regions of cortex (*21, 53*), such connections were not included in the model, which focused on effects within a cortical column. Nonetheless, our results on directly activated neurons are consistent with Histed’s findings (*31*). However, Jankowska and colleagues concluded that the dominant mode of neural activation is indirect (2/3 indirect and 1/3 direct) (*25*), inconsistent with Histed’s findings. This difference might be because Histed et al. investigated neurons very close to the stimulation electrode, where direct activations are likely to occur, instead of a wider space like Jankowska et al., where trans-synaptic activations are more likely.

## Acknowledgements

The authors would like to thank Aman Aberra for helpful discussions on this work.

## Author Contributions

K.K., J.S., L.E.M., S.J.B., and W.M.G. conception and design of research; K.K. implemented computational model and ran simulations; K.K. and J.S. analyzed data; K.K., J.S., L.E.M., S.J.B., and W.M.G. interpreted results; K.K. prepared figures and drafted manuscript; K.K., J.S., L.E.M., S.J.B., and W.M.G. edited and revised manuscript; K.K., J.S., L.E.M., S.J.B., and W.M.G. approved final version of manuscript.

## Supplementary Figures

**Figure S1:**
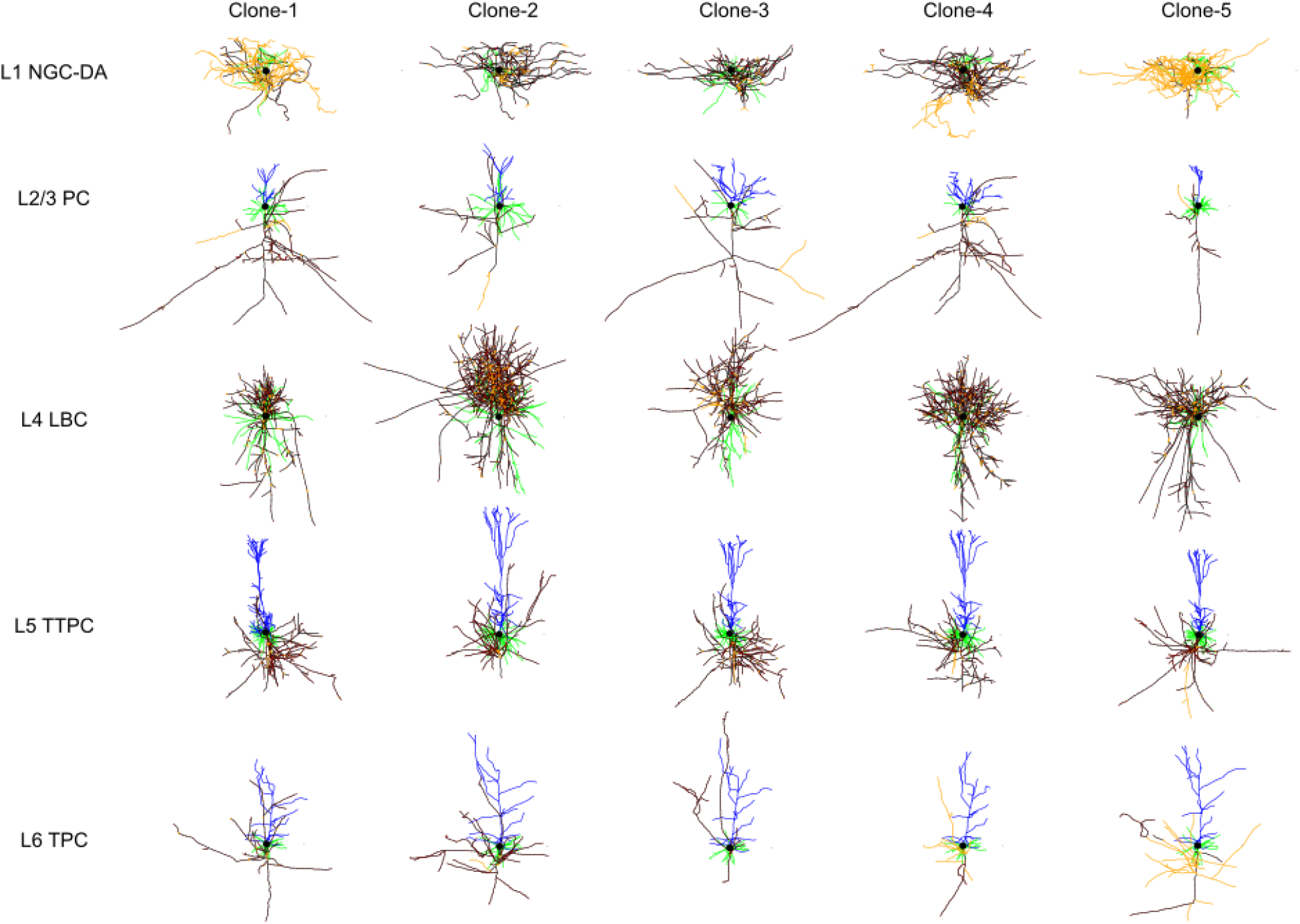
3D morphology of clones of cortical neurons.

**Figure S2:**
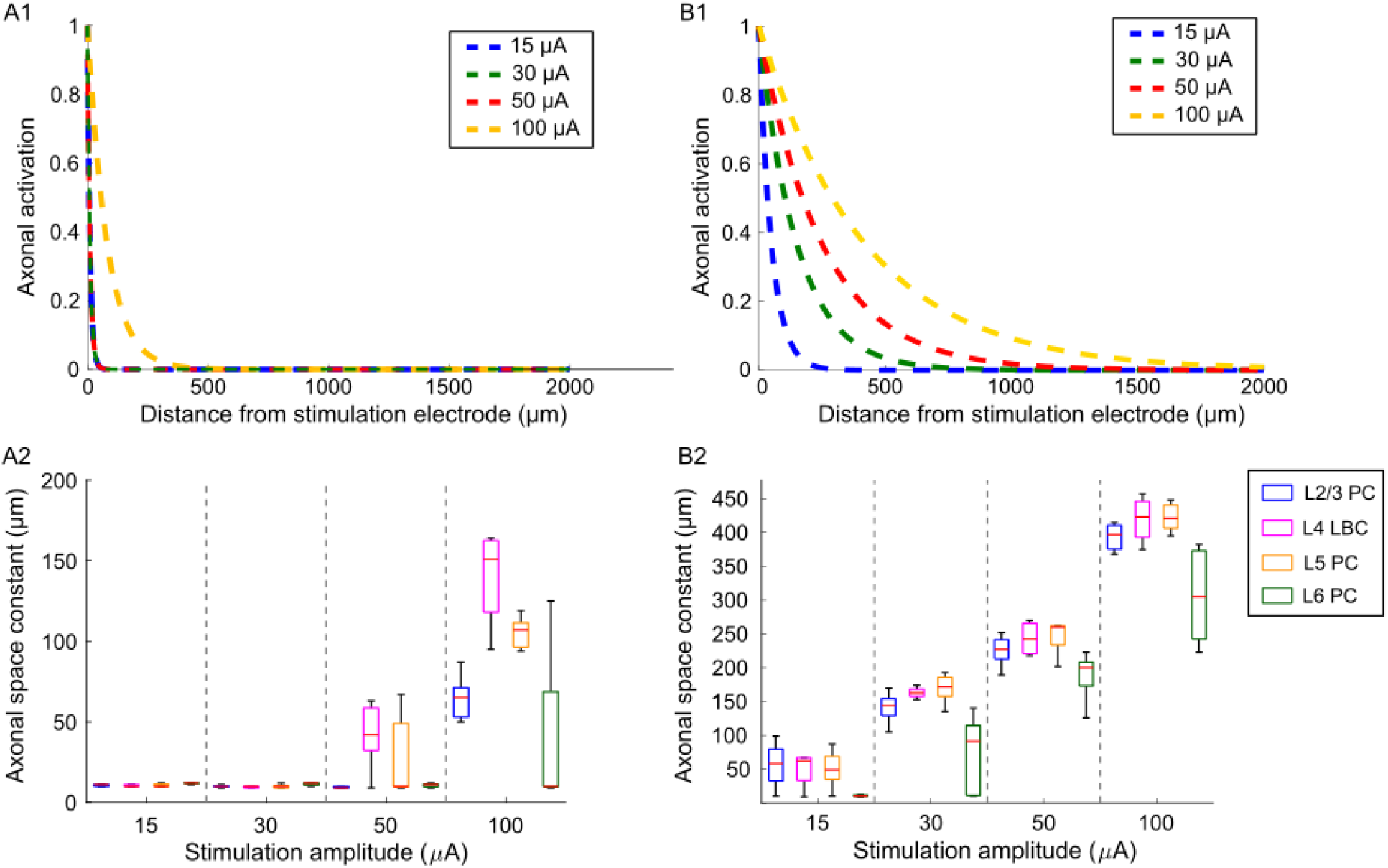
Difference in excitability to ICMS intensity between unmyelinated (*32*) and myelinated (*33*) versions of model neurons. (A1) Activation of unmyelinated L2/3 PC axons as a function of distance from the electrode tip for various ICMS intensities. Responses fall-off exponentially with distance, and the responses are the same for intensities up to 50 μA. (A2) Space constants across cell types are derived from the response shown in A1. (B1) Activation of myelinated L2/3 PC axons as a function of distance from the electrode tip for various ICMS intensities. (B2) Space constants across cell types are derived from the response shown in B1. Space constants increase with stimulus intensity similar to those seen in physiological recordings.

**Figure S3:**
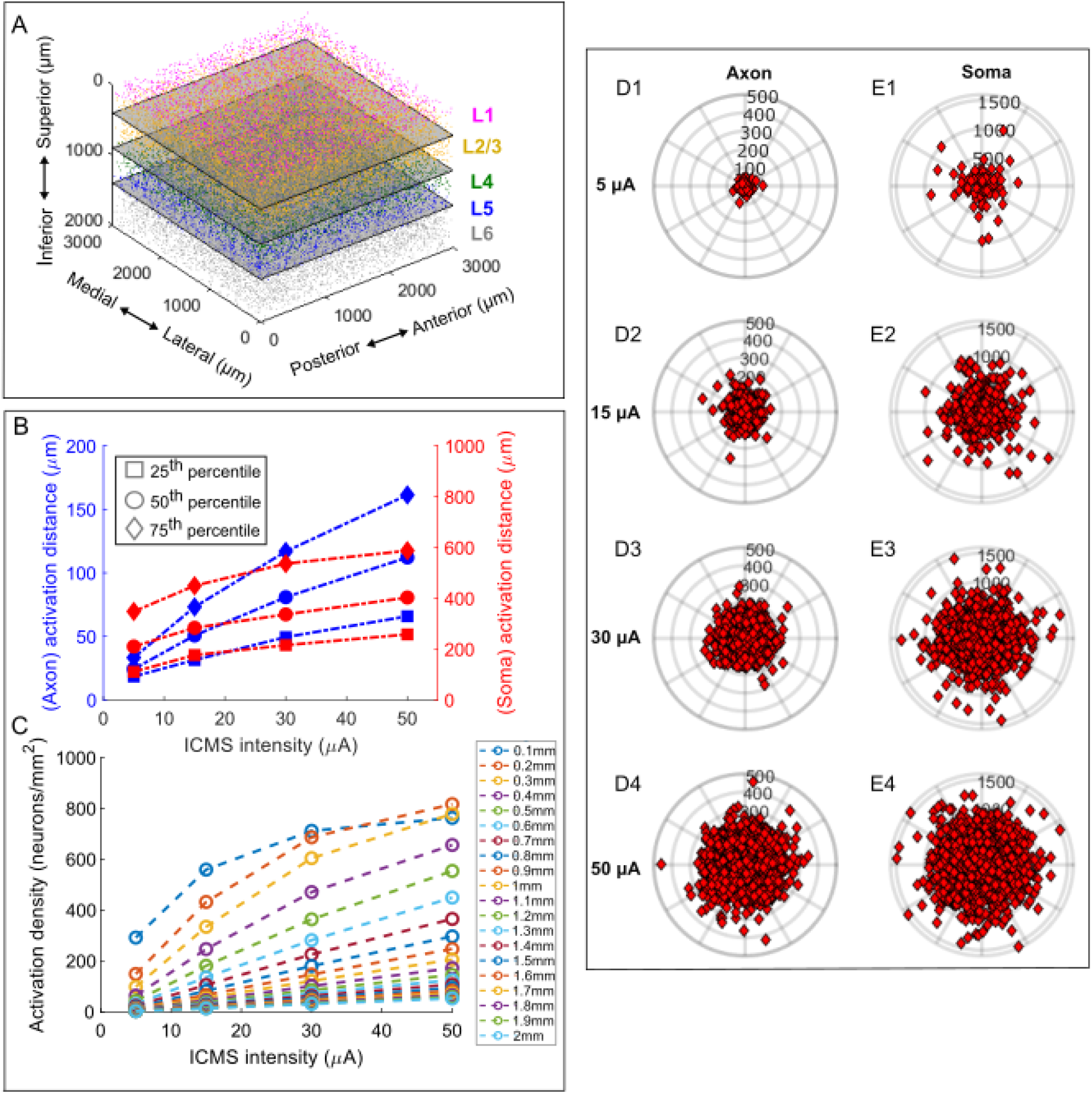
Response to intracortical microstimulation (ICMS) at different intensities. (A) 2D polar plane (axial) was used to visualize the location of activated somas and axons shown in Fig-3A, B. (B) 25^th^, 50^th^ and 75^th^ percentile of activation distance across axons/somas as a function of stimulation intensity. (C) Activation density as a function of ICMS intensity for different radial distances. Polar plots (derived from axial plane) showing the location (radial distance) of activated axon and soma to ICMS at (D1, E1) 5 μA, (D2, E2) 15 μA, (D3, E3) 30 μA and (D4, E4) 50 μA. Theta for each neuron was randomly assigned from a uniform distribution between 0-360 degrees.

**Figure S4:**
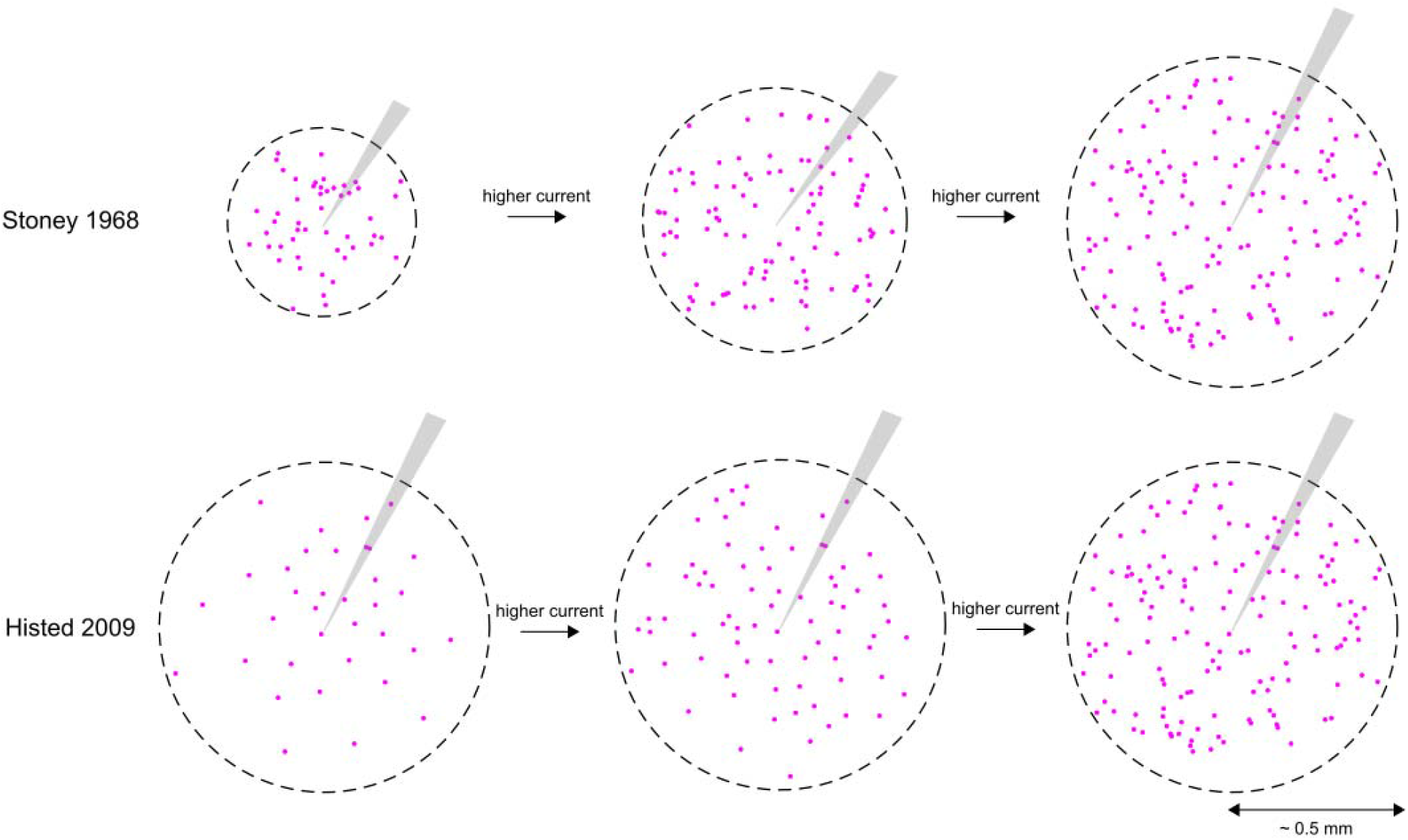
Two models on the effect of ICMS intensity on the radial distance of somatic activation. (A) Stoney et al., 1968 proposed radial distance of activation to increase with stimulation intensity based on the equation. (B) Histed et al., 2009 proposed the radial distance of activation does not increase with stimulation intensity, whereas an increase in intensity results in an increase in the number of activated neurons. Our model-based results confirm Histed et al., 2009 findings on the effect of stimulation intensity on radial distance of somatic activation, and are consistent with the Stoney et al. 1968 model for axonal activation.

